# Illuminating kinase inhibitors biology by cell signaling profiling

**DOI:** 10.1101/2022.04.03.486887

**Authors:** Alexander V. Medvedev, Sergei Makarov, Lyubov A. Medvedeva, Elena Martsen, Kristen L. Gorman, Benjamin Lin, Sergei S. Makarov

## Abstract

Protein kinase inhibitors (PKI) are promising drug candidates for many diseases. However, even selective PKIs interact with multiple kinases and non-kinase targets. Existing technologies detect these interactions but not the resultant biological effects. Here, we describe an orthogonal PKI evaluation approach that entails fingerprinting of cell signaling responses. As the readout, we profiled the activity of 45 transcription factors linking signaling pathways to genes. We found that inhibitors of the same kinase family exhibited a consensus TF activity profile (TFAP) invariant to PKI chemistry and mode of action (allosteric, ATP-competitive, or genetic). Specific PKI consensus signatures were found for multiple kinase families (Akt, CDK, Aurora, RAF, MEK, and ERK) with high-similarity consensus signatures of signaling cascade kinases. Thus, the PKI consensus signatures provide bona fide markers of cell response to on-target PKI activity. However, the consensus signatures appeared only at certain inhibitor concentrations (‘on-target windows’). Using concentration-response signature analysis, we identified PKI interactions dominating cell response at other concentrations. Finally, we illustrate this approach by selecting putative chemical probes for evaluated kinases. Therefore, the effect-based TFAP approach illuminates PKI biology invisible to target-based technologies and provides clear quantitative metrics to aid the selection of polypharmacological PKIs as chemical probes and drug leads.

## INTRODUCTION

Protein kinases are considered the most promising drug targets for cancer, and kinase drug discovery has expanded into immunological, inflammatory, degenerative, metabolic, cardiovascular, and infectious diseases(*1*). One of the most challenging issues is PKI promiscuity stemming from the homology of active kinase centers(*2*). Selectivity profiling studies showed that even the most selective PKIs inhibit multiple kinases(*3*–*5*). Moreover, like other drugs, PKIs interact with various non-kinase targets, such as bromodomain-containing proteins, G protein-coupled receptors, cytoskeleton, prostaglandin synthases, AhR, ferrochelatase, and NQO(*6*).

PKI polypharmacology poses difficult problems to drug developers. On the one hand, it can cause undesirable adverse effects. On the other hand, multi-kinase inhibitors proved highly effective for cancer treatment(*1, 7*). Polypharmacology is also a challenging issue for selecting PKI chemical probes for examining the biological function of kinases and validating them as drug targets(*8, 9*).

The most widely used evaluation strategy aims to detect PKI targets by various biochemical and cell-based assays and proteomics techniques(*3, 10, 11*). The resulting datasets characterize PKI affinity, activity, and target occupancy across kinases and non-kinase proteins. However, these data do not predict the cumulative biological outcome of polypharmacological inhibitors and do not inform which PKI interactions dominate cells’ response at a given concentration. Answering these questions requires alternative, effect-based approaches.

Here, we describe an orthogonal, effect-based PKI evaluation approach based on fingerprinting of cellular signal responses. Previously, we showed that analysis of cell signaling responses to various perturbagens permitted pinpointing the targeted biological processes and cell systems(*12*). As kinases regulate critical nodes in signal transduction, we expected to find such patterns for kinase inhibitors.

To capture cell signaling responses, we profiled the activity of transcription factors (TFs) that lie at the apexes of signaling pathways and connect them to regulated genes. TFs are a class of proteins that bind specific gene sequences, thereby modulating transcription. The human genome has an estimated 1,600 TFs comprising a few hundred TF families with similar binding sequences(*13*). To capture multiple TF responses, we used our multiplexed reporter system (the FACTORIAL®) that enables quantitative parallel TF activity assessment in a single well of cells(*14*).

The FACTORIAL generates highly reproducible cell response signatures termed TF activity profiles (TFAP). These signatures are quantitative descriptors that allow comparing cells’ responses to compounds using straightforward correlation analysis.

Here, we assessed inhibitors of six kinase families with diverse biological functions. One of those was the PKB/Akt family that regulates multiple cell functions, including glucose metabolism, apoptosis, cell proliferation, and migration(*15*). Another family comprised cyclin-dependent kinases (CDK) that regulate cell cycle and transcription(*16*). We also evaluated inhibitors of Aurora kinases that regulate chromatid segregation in cell division(*17*). In addition, we examined inhibitors of RAF, MEK, and ERK kinases constituting the ERK MAPK cascade that plays pivotal role in cell proliferation, differentiation, and oncogenic transformation(*18*). The evaluated PKI panel included inhibitors with different mode of action (MOA) and chemical structures.

The main finding is that inhibitors of a given kinase family at some concentrations exhibit a consensus TFAP signature invariant to PKI chemistry and MOA. We show that each evaluated kinase family has a specific PKI consensus signature. Thus, the PKI consensus signatures provide bona fide markers of cell response to on-target PKI activity. Further supporting this notion, functionally related ERK MAPK cascade kinases had virtually identical PKI consensus signatures.

However, individual inhibitors elicited the ‘on-target’ PKI consensus signatures only within certain concentration ranges that we termed ‘on-target windows’. Concentration-response signature analysis permitted determining which PKI interactions dominated cells’ response at other concentrations. To illustrate applications of this approach, we selected PKIs with the largest on-target windows as putative chemical probes. Therefore, the TFAP approach illuminates PKI biology invisible to target-based approaches and complements existing techniques by clear quantitative metrics to aid the selection of PKI drug leads and chemical probes.

## RESULTS

### Fingerprinting TF responses by the FACTORIAL

Cellular signaling pathways regulate TF activity through various posttranslational modifications, e.g., phosphorylation, acetylation, etc., that do not alter TF protein content. Hence, TF activity assessments require functional assays, such as DNA-binding and reporter gene assays(*19*). However, existing assays are not well suited for quantitative profiling of multiple TFs’ activity.

Therefore, in this study we used a multiplexed reporter system, the FACTORIAL(*14*). It comprises a set of TF-specific reporter plasmids termed reporter transcription units (RTUs) that are similar to traditional reporter gene constructs, but RTU activity is evaluated by detecting reporter RNA transcripts.

A FACTORIAL assay entails transient co-transfection of RTU plasmids into assay cells followed by reporter RNA profiling. The FACTORIAL is built on a homogeneous design ensuring equal detection efficacy across RTUs (Fig. S1). This design minimizes experimental variability and provides outstanding reproducibility and the signal/noise ratio(*14*). As a result, the assay can detect with high fidelity even weak TF responses(*12*).

The TFAP signature endpoints characterize the activity of 45 TF families and two minimal promoters (listed in Fig. S1), thus capturing essential TF responses to physiological and pathological stimuli, including xenobiotics, growth factors, inflammation, and various stress factors(*14*).

### Capturing TFAP signatures of kinase inhibitors

This study examined inhibitors of Akt, cyclin-dependent kinases (CDK), Aurora, RAF, MEK, and ERK kinase families. As the enabling technology, we used extensively validated FACTORIAL assay in human hepatocyte HepG2 cell line(*12, 14, 20*).

As in previous studies, we assessed TFAP signatures after a prolonged (24 h) treatment to obviate differences in PKI pharmacokinetics and rapid transitory responses. Previously, we determined that a prolonged exposure produced stable perturbagens’ signatures reflecting a steady activation state of cell signaling(*12*).

The PKI TFAP signatures are presented as radial graphs with log axes showing TF activity fold changes in PKI-treated vs. vehicle-treated cells. By this definition, the baseline TFAP signature of vehicle-treated cells is a perfect circle with R=1.0 (the “null” signature). Presented here TFAP signatures are the average of at least three independent FACTORIAL assays.

We estimated the similarity of cell responses to compounds as the Pearson correlation of TFAP signatures. We set the identity threshold at r=0.70, since the probability of two random 47-endpoint signatures correlating with r> 0.70 is very low (*P*<10^−7^)(*21*). To find a consensus signature among multiple TFAP signatures, we used unsupervised cluster analysis(*12*).

### The on-target TFAP signatures of kinase inhibitors

#### The consensus signature of AKT inhibitors

The Akt family comprises Akt1, Akt2, and Akt3 kinases that respond to various stresses, hormonal and nutrition factors to regulate multiple biological functions(*15*). The PKI set comprised four ATP-competitive pan-Akt inhibitors and one allosteric Akt 2/3 inhibitor (Fig. 1A). In addition, we used a genetic approach comprising ectopic expression of a dominant negative (DN) kinase-dead Akt (K179M) cDNA. Each inhibitor was evaluated at multiple concentrations.

**Fig. 1.**
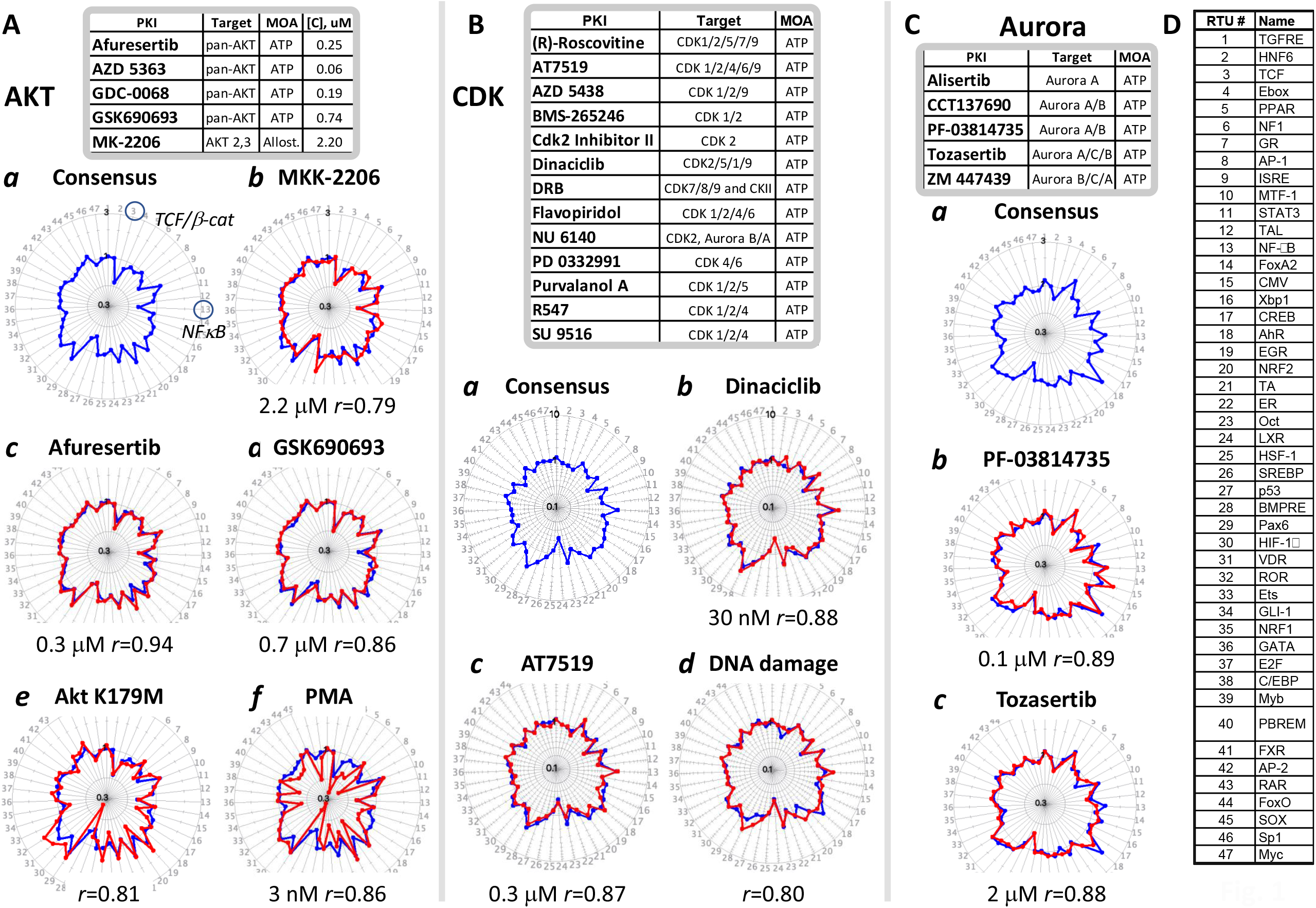
The TFAP signatures of Akt, CDK, and Aurora inhibitors. The TFAP signatures were assessed by FACTORIAL assay in human hepatocyte HepG2 cells after a 24-h treatment with indicated inhibitors. The consensus signatures were obtained by clustering signatures of individual PKIs at multiple concentrations. (**A)** The consensus signature of Akt inhibitors (blue) (**a**) overlaid by signatures of indicated Akt inhibitors (red) (**b-f**). (**B)** The consensus signature of CDK inhibitors (**a**) (blue) overlaid by signatures of indicated CDK inhibitors (red) (**b-e**). (B**d)** Overlayed consensus signatures of CDK inhibitors (blue) and DNA damaging agents (red). (**C)** The consensus signature of Aurora inhibitors (**a**) (blue) overlaid by individual inhibitors’ signatures (**b**,**c**) (red). **D**. The TFAP signature endpoints (detailed endpoints’ description in Fig. S1). The similarity of consensus and individual PKI signatures is Pearson correlation coefficient ***r***. Each TFAP signature is average data of at least three independent FACTORIAL assays. The signature axes show fold-changes of TF activity in PKI-treated vs. vehicle-treated cells on a log scale from 0.3 to 3.0 (**A, C**) and 0.1 to 10.0 (**B**).

Clustering the obtained signatures showed that all tested Akt inhibitors at certain concentrations elicited a consensus TFAP signature (Fig. 1Aa). This consensus signature comprised comparatively weak (≤two-fold) TF responses; nonetheless, the consensus signature perfectly matched individual inhibitors’ signatures (Figs. 1Ab-e) (see also Fig.4 for concentration-response analysis of Akt inhibitors’ signatures). Importantly, TF responses of the consensus signature were in agreement with known Akt TF targets, such as TCF/β-catenin(*22*) and NFκB(*23*) (Fig. 1Aa).

Therefore, different Akt inhibitors at certain concentrations produced identical TFAP signatures, regardless of their chemistry, MOA, and specific kinase targets within the Akt family. That suggests that the consensus PKI signature provides a bona fide marker for on-target Akt PKI activity. Further supporting this notion, the Akt PKI consensus signature matched the signature elicited by a prototypical protein kinase C activator phorbol 12-myristate 13-acetate (PMA) (Fig. 1Af). This observation is consistent with established role of PKC as a negative regulator of Akt kinase(*24*).

#### The consensus signature of CDK inhibitors

The CDK family comprises 21 kinases with diverse cellular functions. CDK1, −2, −4 and −6 play key roles in the cell cycle regulation, whereas CDK8–9 and −19 regulate gene transcription, and CDK7 has broader roles in both processes(*16*). CDK activity is modulated by interactions with cyclins and CDK inhibitory proteins and by posttranslational modifications(*16*).

Here, we evaluated a panel of 13 ATP-competitive inhibitors with different selectivity profiles across the CDK family (Fig. 1B). We found a consensus TFAP signature (Fig 1Ba) elicited at certain concentrations by the tested inhibitors (Fig. 1Bb-c). (for more CDK PKI signatures, see Fig. S2). Querying this on-target consensus signature against our database of landmark perturbagens retrieved the consensus signature of cell response to high concentrations of DNA damaging agents (auramine, camptothecin, cisplatin, and UV radiation)(*12*) (Fig. 1Bd). Therefore, CDK inhibitors with different chemical structures and selectivity had a distinct consensus TFAP signature plausibly reflecting the pivotal role of this kinase family in DNA damage response(*25*).

#### The consensus signature of Aurora inhibitors

The Aurora family comprises Aurora A, B, and C kinases that regulate chromatid segregation in cell division(*17*). The evaluation of five Aurora ATP-competitive inhibitors (Fig. 1C) revealed a consensus TFAP signature (Fig. 1Ca) with high similarity to Aurora inhibitors with different selectivity profiles (Figs. 1Cb-c and S3).

#### Consensus signatures of RAF, MEK, and ERK inhibitors

Many cellular kinases are integrated into kinase cascades that amplify and relay regulatory signals from the cell periphery to the nucleus. An example is mitogen-activated protein kinase (MAPK) cascades that regulate proliferation, differentiation, apoptosis, and stress responses(*26*). One of those, ERK MAPK cascade comprises RAF, MEK, and ERK kinases(*18*). This cascade is regulated by the small guanine nucleotide-binding protein Ras that activates RAF kinase, which phosphorylates and activates MEK1 and MEK2 kinases. MEK kinases, in turn, phosphorylate and activate the most distal ERK1 and ERK2 kinases(*27*). The Ras/RAF/MEK/ERK cascade is critically involved in oncogenic transformation, as evidenced by mutational activation of its components in cancer, including the most frequently mutated in human cancers Ras oncogene(*28*).

The evaluation of five ATP-competitive pan-RAF inhibitors and two B-RAF inhibitors (Fig. 2A) revealed consensus TFAP signature (Fig. 2Aa) elicited at certain concentrations by all these inhibitors (Fig. 2Ab-c) (more RAF PKI signatures shown in Fig. S4).

**Fig. 2.**
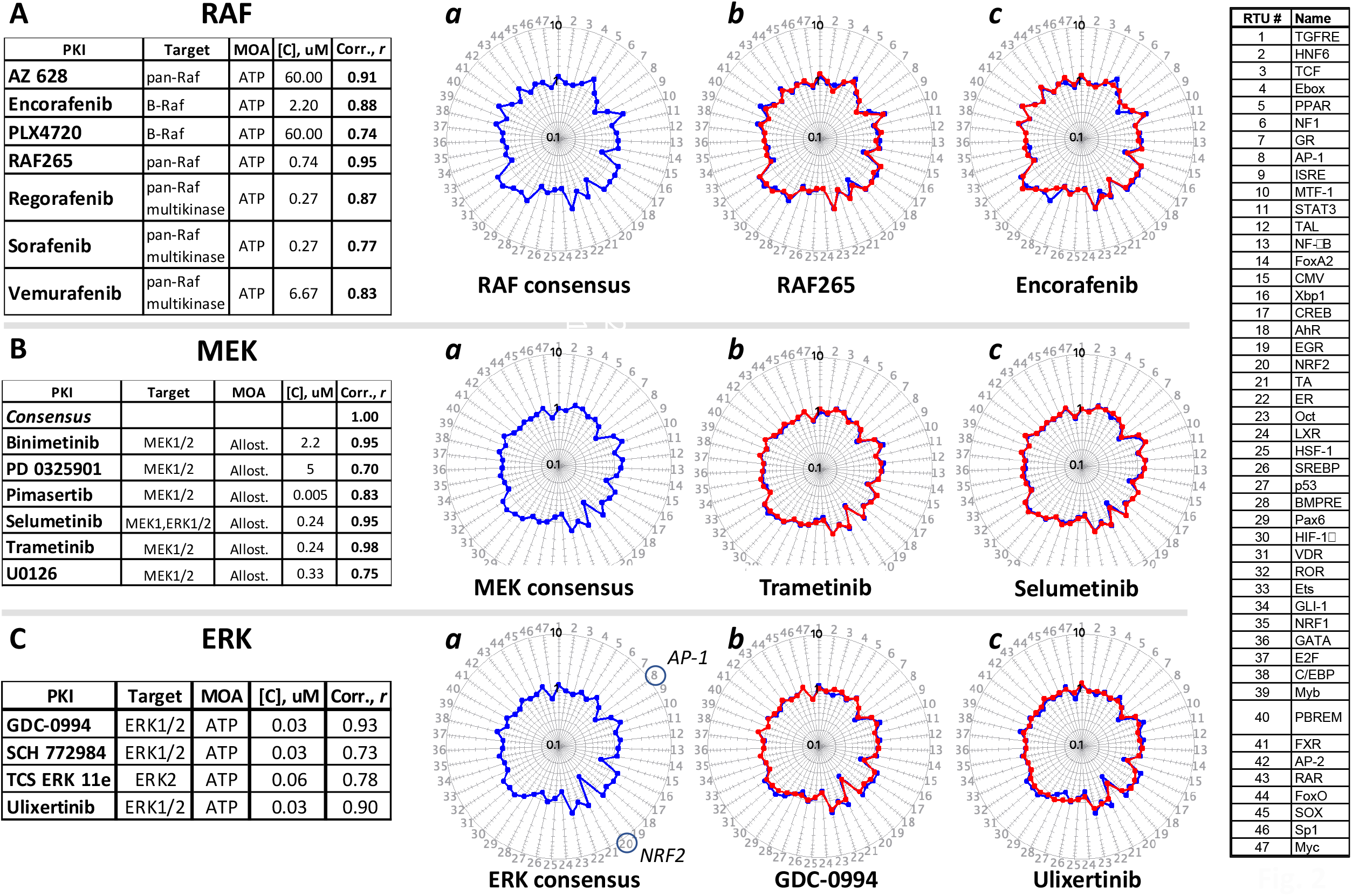
The TFAP signatures of RAF (A), MEK (B), and ERK (C) inhibitors. The TFAP signatures were analyzed as described by Fig. 1 legend. The signatures’ axes show fold-changes of TF activity in PKI-treated vs. vehicle-treated cells on a log scale from 0.1 to 10.0.

Similar to that, we found consensus TFAP signature for six allosteric MEK1/2 inhibitors (Fig. 2Ba-c) (more signatures of MEK inhibitors are shown in Fig. S5). Likewise, ERK1/2 a panel of ERK inhibitors (Fig. 2C) also had a distinct consensus signature (Figs. 2C and S6).

### Inhibitors of signaling cascade kinases have high-similarity on-target signatures

Further analysis showed that PKI consensus signatures of RAF, MEK, and ERK kinases had remarkably high pairwise similarity values (r≥0.79), exceeding the identity threshold (Fig. 3). Notably, individual TF responses of these consensus signatures agreed with known TF targets of this kinase cascade, e.g., AP-1(*18, 28*) and NRF2(*29*) (Fig. 2Ca). By contrast, we found little similarity of the PKI consensus signatures of CDK, Akt, and Aurora kinases to each other and the RAF/MEK/ERK kinases (Fig. 3). Therefore, the similarity of PKI consensus signatures reflects the functional relationship of corresponding kinases.

**Fig. 3.**
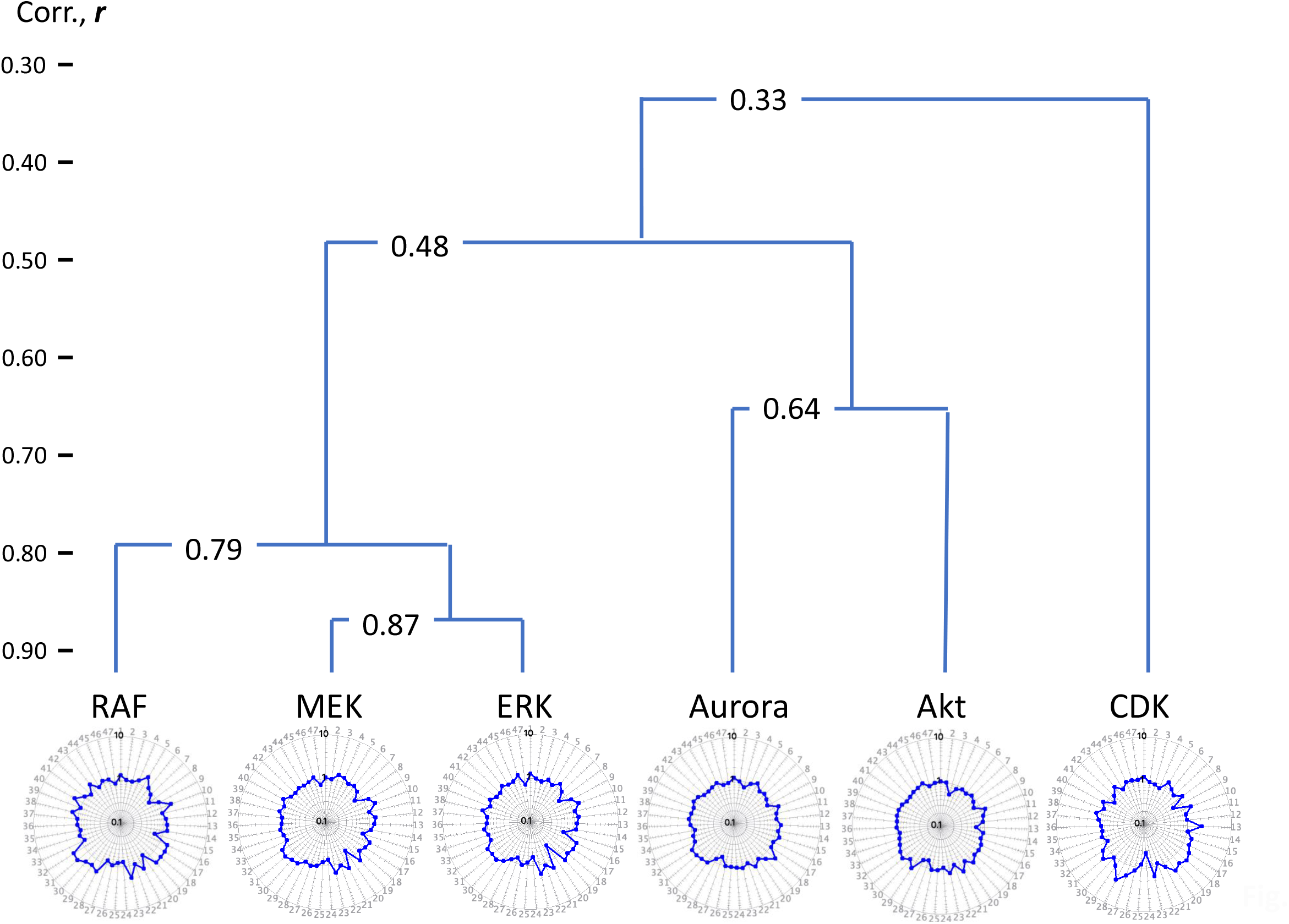
Signaling cascade kinases have similar PKI consensus signatures. The signature tree shows PKI consensus signatures of Figs. 1-2. The similarity was calculated by agglomerative cluster analysis as Pearson correlation ***r***. The TF activity fold-changes for all signatures is shown on a log scale from 0.1 to 10.0.

### Assessing off-target PKI activity by concentration-response signature analysis

The PKI evaluation showed that thier on-target PKI consensus signatures appeared only within certain concentration ranges that we termed the ‘on-target windows’. At other concentrations, we observed different signatures, presumably reflecting off-target responses to PKIs. To ascertain the underlying PKI activity, we queried our dataset of landmark perturbagens and the PKI consensus signatures. The following data (Figs. 4-7) illustrate this approach.

**Fig. 4.**
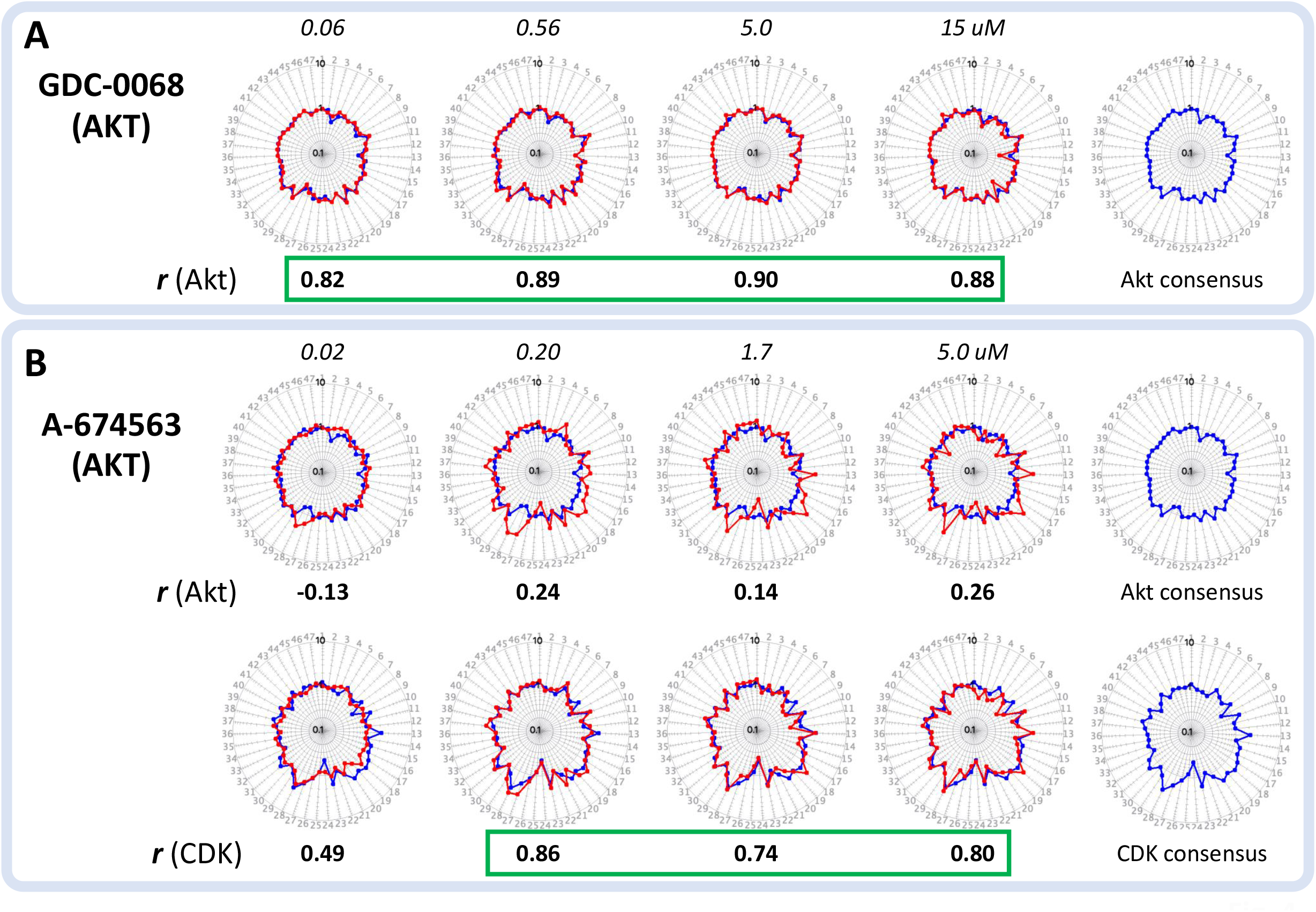
The concentration-response signature analysis of AKT inhibitors. The signatures were assessed as described in Fig. 1 legend. Signatures’ axes show fold-changes of TF activity in PKI-treated vs. vehicle-treated cells on a log scale from 0.1 to 10.0. **A**. Signatures of GDC 0068 (red) overlayed by the consensus signature of AKT inhibitors (blue). **B**. Signatures of A 674563 (red) overlayed by the consensus signature of AKT (upper panel) or CDK (bottom panel) inhibitors. The overlayed signatures’ similarity is Pearson correlation ***r***. Signatures with r values exceeding the identity threshold (r>0.70) are outlined. Each TFAP signature is the average data of at least three independent FACTORIAL assays.

#### Akt inhibitors

Fig. 4 shows concentration-response signatures of two Akt inhibitors. One was a pan-Akt inhibitor GDC-0068 with IC50 for of Akt1/2/3 of 5 nM/18 nM/8 nM in cell-free assays(*30*). The other inhibitor was A674563, a multi-kinase inhibitor not included into the panel of Akt inhibitors of Fig. 1A. In addition to Akt1 (IC50 of 11 nM), A674563 inhibits PKA and Cdk2 (16 nM and 46 nM)(*31*), and FLT3 (2.1 μM)(*32*).

GDC-0068 signatures matched the consensus Akt PKI signature at all tested concentrations (60 nM to 15 μM) (Fig. 4A). In contrast, A674563 signatures had a low similarity to the Akt PKI consensus signature (Fig.4B, upper panel), instead matching the consensus signature of CDK inhibitors (Fig. 4B, lower panel). Therefore, cells’ response to GDC-0068 was dominated by Akt inhibition. However the response to A674563 was primarily determined by CDK inhibition, even though its reported IC50 value for Akt (11 nM) was lower than for CDK (46 nM).

#### MEK/ERK inhibitors

Fig. 5A shows TFAP signatures for non-ATP-competitive allosteric MEK1/2 inhibitors Selumetinib (AZD6244) (Aa) and Pimasertib (AS-703026) (Ab). Selumetinib’s signatures perfectly matched the consensus signature of MEK inhibitors at concentrations up to 60 μM. In contrast, Pimasertib’s signatures were identical to the consensus signature at lower concentrations but, starting with 0.27 μM, it showed off-target activity at the aryl hydrocarbon receptor (AhR). A similar off-target activity at AhR was also found for MEK inhibitor, U0126 (Fig. 5Ac), which agreed with data by others(*33*).

**Fig. 5.**
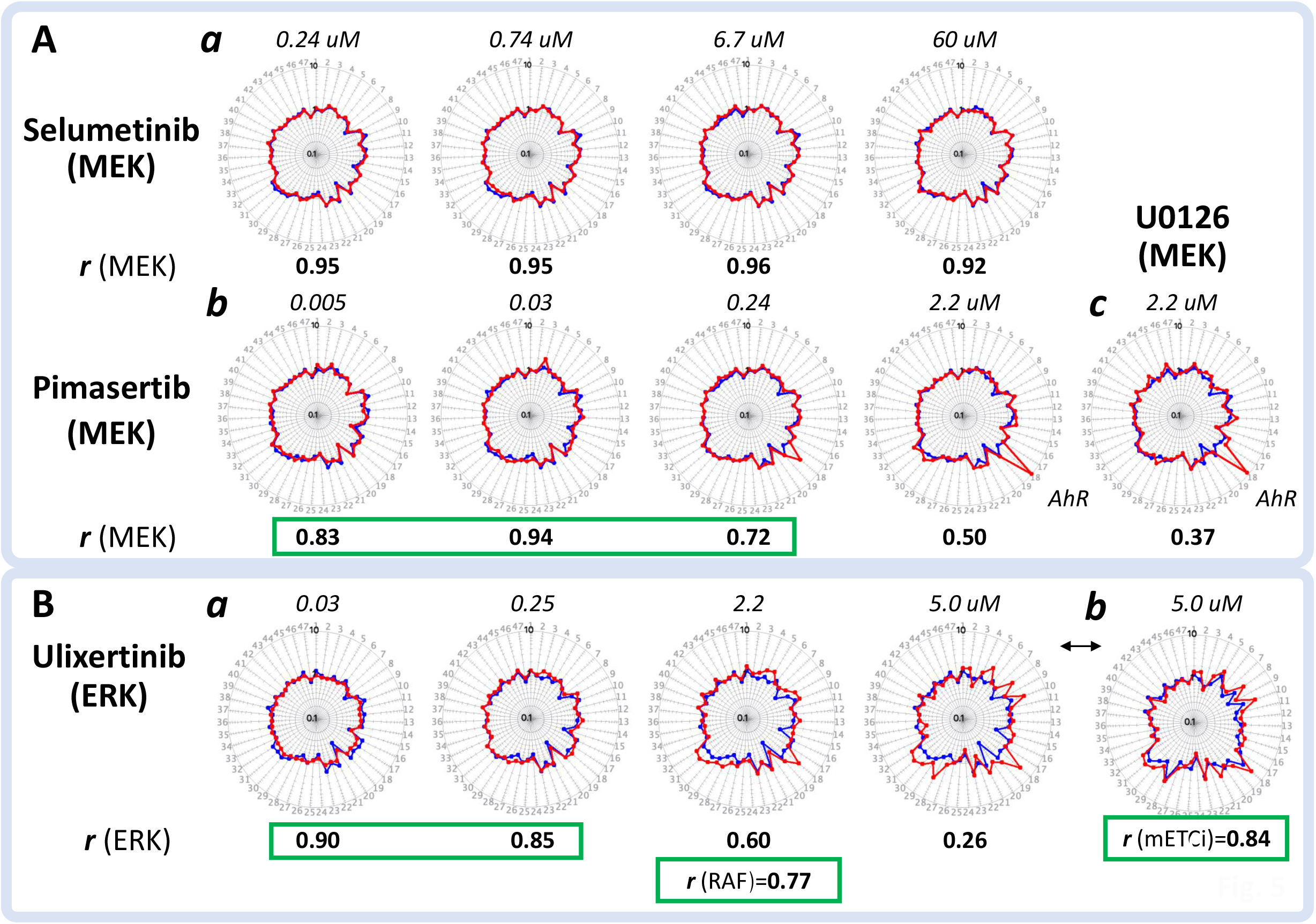
The concentration-response signature analysis of MEK and ERK inhibitors. **A**. Signatures of Selumetinib (**a**), Pimaserib (**b**), and U 0126 (**c**) (red) overlayed by the consensus signature of MEK inhibitors (blue). **B**. Signatures of Ulixertinib (in red) overlayed by the consensus signatures of ERK inhibitors (**a**) or mitochondria inhibitors (mETCi) (**b**) (blue). The overlaid signatures’ similarity is the Pearson correlation ***r***. Each TFAP signature is the average data of at least three independent FACTORIAL assays. Identical signatures (r>0.70) are outlined.

Fig. 5B shows TFAP signatures of ERK1/2 inhibitor Ulixertinib (BVD-523)(*34, 35*). Its signatures were identical to the consensus signature of ERK inhibitors at concentrations from 30 nM and 250 nM. However, the signature at 2.2 μM best matched the consensus signature of RAF inhibitors, and the best match at 5 μM was the consensus signature of mitochondria inhibitors (Fig. 5Bb) that was described in our previous report(*12*). The inferred mitochondria inhibiting activity was confirmed by a cell-based functional assay (Fig. S7).

#### CDK inhibitors

Fig. 6 shows signatures of Dinaciclib and NU6140. Dinaciclib is a CDK2, CDK5, CDK1, and CDK9 inhibitor(*36*), whereas NU6140 was developed as a selective CDK2 inhibitor(*37*).

**Fig. 6.**
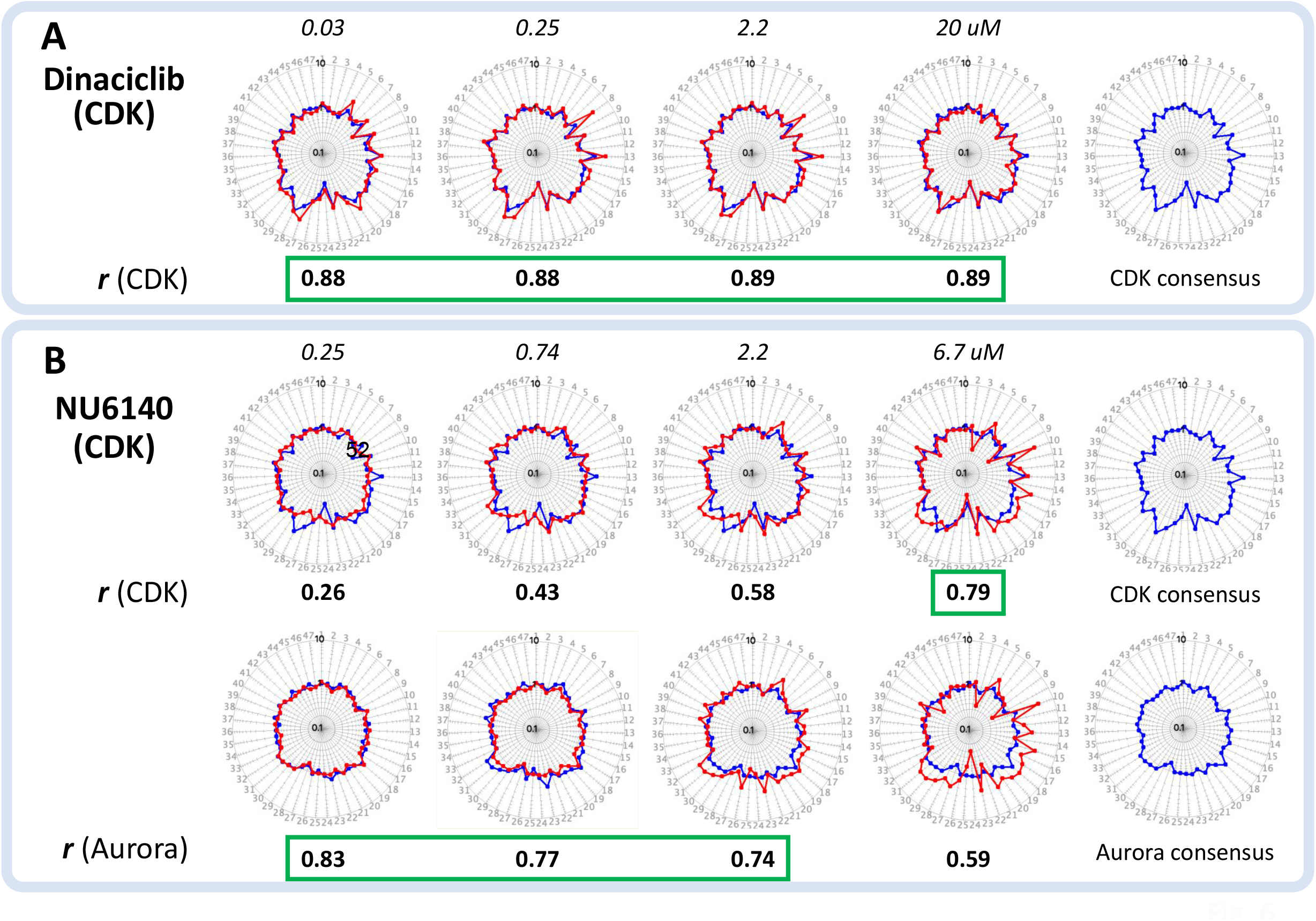
The concentration-response analysis of CDK inhibitors. **A**. Dinaciclib’s Signatures at indicated concentrations (red) overlayed on the consensus signature of CDK inhibitors (blue). **B**. NU 6140 signatures (red) overlayed on the consensus signatures of CDK (upper panel) or Aurora (bottom panel) inhibitors. The ‘similarity of overlayed signatures is Pearson correlation ***r***. Each signature is the average data of at least three independent FACTORIAL assays. Identical signatures (r>0.70) are outlined.

Dinaciclib’s signatures were identical (r>0.70) to the consensus signature of CDK inhibitors at all tested concentrations (Fig. 6A). In contrast, NU6140 signatures matched the CDK PKI consensus signature only at a high concentration (6.7 μM) (Fig. 6B, upper panel). At lower concentrations, NU6140 signatures were identical to the consensus signature of Aurora inhibitors (Fig. 6B, lower panel). These results are consistent with the data on inhibition of Aurora A/B kinases at lower NU6140 concentrations (IC50 of 67 and 35 nM, respectively)(*38*) vs. CDK2 (0.4 μM)(*37*).

### Guiding chemical probe selection by TFAP signatures

Chemical probes are selective small-molecule modulators of kinase activity. These probes provide indispensable tools for examining the biological functions of kinases in cells and animals and validating kinase drug targets(*9, 39*). The most essential requirement is that cells’ response must be predominantly determined by probe’s activity toward the intended kinase. Other important criteria include defined working range for the specific probe activity, the bioavailability, and chemical stability in the evaluated biological system(*8, 40, 41*).

Here, we used the concentration-response analysis of PKI signatures to select putative chemical probes. We searched for inhibitors exhibiting the on-target PKI consensus signature in the broadest concentration ranges, with the assumption that cells’ response at these concentrations is dominated by the on-target PKI activity. Importantly, as we obtained the PKI signatures after a prolonged treatment, this approach also accounts for such essential chemical probe parameters as bioavailability and stability.

Fig. 7 illustrates selecting chemical probes for Akt and MEK kinases.

**Fig. 7.**
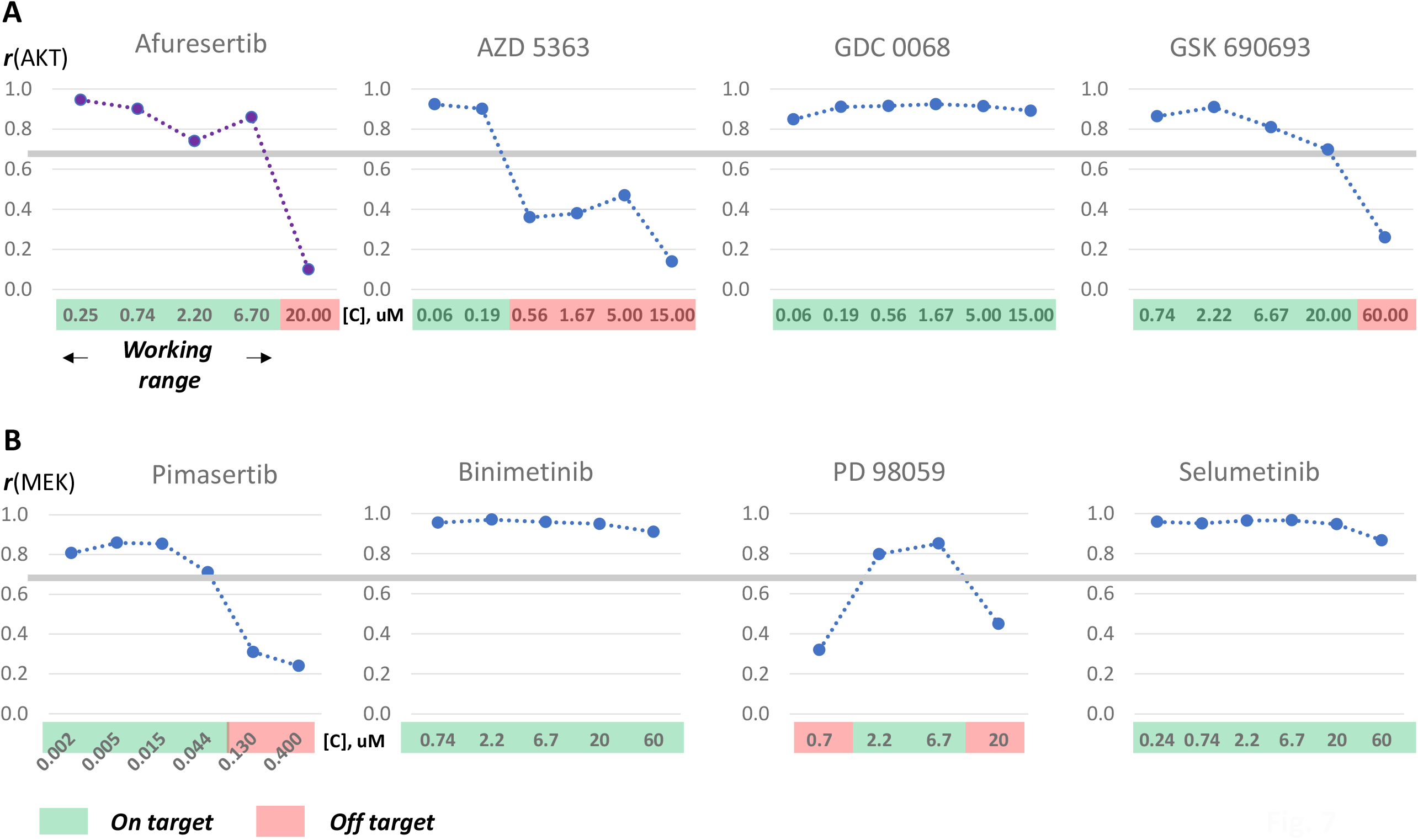
Selecting PKI chemical probes. The similarity values of individual PKI signatures at indicated concentrations vs. the consensus signatures of Akt (**A**) and MEK (**B**) inhibitors. Working ranges (green) are inhibitors’ concentrations eliciting the on-target consensus signatures r≥0.70 (grey line). Each TFAP signature is an average of at least three independent FACTORIAL assays.

#### Akt

The largest on-target window (from 60 nM to 60 μM) showed **GDC-0068** (Fig. 7A).

#### MEK

The largest on-target windows were found for **Binimetinib (**0.74 to 60 μM) and **Selumetinib** (0.24 to 60 μM) (Fig. 7B). Another putative MEK probe is **Trametinib** (0.24 to 60 μM) (Fig. S5).

Based on the concentration-response analysis, the putative probes for other kinase families are (working range in parentheses):

*RAF:* **RAF265** (80 nM to 60 μM) and **Encorafenib** (0.74 to 60 μM) (Fig. S4);

*ERK:* **GDC0994** (9nM to 60 μM)(Fig. S6);

*CDK:* **Dinaciclib** (30 nM to 60 μM) and **Flavopiridol** (70 nM to 20 μM) (Fig. S2);

*Aurora:* **PF-03814735** (10 nM to 3 μM) and **ZM 447439** (0.25 to 20 μM) (Fig. S3).

## DISCUSSION

PKI promiscuity is a major challenge to kinase drug development. The existing paradigm predicates PKI activity on its target interactions, assuming a predominant role for interactions with the lowest IC50. However, elucidating the cumulative biological effect of multiple interactions requires additional effect-based approaches.

Here, we described an orthogonal evaluation approach by assessing PKI-induced cell signaling responses and using TF activity as the readout. Previously, we showed that perturbagens of various biological processes and cell systems produced characteristic TF activity profiles(*12*). For example, specific TFAP signatures were found for mitochondria inhibitors, cytoskeleton disruptors, DNA damaging agents, and inhibitors of Ub/PS protein degradation and histone deacetylation(*12*). As kinases lie at the heart of cell signaling, here we applied the TFAP approach to PKI evaluation. The main findings of this paper are:

1. cells’ exposure to PKIs induces distinct and reproducible TF activity profiles;
2. inhibitors of the same kinase family at certain concentrations exhibit a consensus TFAP signature invariant to PKI chemistry and MOA;
3. multiple kinase families with diverse biological functions have specific PKI consensus signatures;
4. functionally related signaling cascade kinases have similar PKI consensus signatures, and
5. polypharmacological PKIs exhibit multiple signatures at different concentrations. Their analysis reveals the PKI interactions playing a dominant role at a given concentration.

Consistent with our previous report(*12*), these findings indicate that kinase inhibitors, too, cause coordinated cell signaling responses. Furthermore, the PKI consensus signatures provide bona fide markers for the on-target PKI activity. A distinct advantage of TFAP signatures is that these reproducible quantitative descriptors are amenable to straightforward correlation analysis, which obviates the need for probabilistic predictions and complex bioinformatic computations.

The TFAP analysis provided valuable insights into PKI polypharmacology. We showed that, in addition to the consensus signature, most PKIs produced multiple signatures at different concentrations, reflecting their on-target and off-target activity. Querying these signatures against landmark perturbagens permitted identifying the underlying activity. Interestingly, some annotated specific inhibitors did not exhibit the corresponding on-target PKI consensus signature altogether, suggesting a prevailing effect at another kinase (e.g., Fig. 4B). In other cases, cells’ response was dominated by inhibition of different kinases at different PKI concentrations (Fig. 6B). That raises interesting questions about kinases’ hierarchy in cell response.

Our approach also permitted detecting off-target PKI effects at non-kinase targets, such as the AhR activation (Fig. 5A) and mitochondrial malfunction (Fig. 5B). These off-target effects can be detrimental in one clinical settings (*42, 43*) but beneficial in others (*44, 45*). Therefore, the TFAP approach provides clear quantitative metrics for PKI evaluation, including the on-target and off-target signatures and the concentration ranges wherein these activities dominate. These metrics can aid in selecting PKIs for specific therapeutic applications. Furthermore, the TFAP approach can assist in preclinical drug development. For example, the TFAP signatures can guide PKI lead optimization by assessing the biological effects of PKI modifications on the on-target and off-target windows and the appearance of additional off-target responses.

Another important application area of our approach relates to chemical probe selection. We illustrated these applications by selecting PKIs with the largest on-target windows as putative chemical probes. Notably, some of our selections (e.g., Selumetinib) agreed with the expert opinion of the Chemical Probes Portal(*46*). Although these data relate to a particular cell line (HepG2), this approach can serve as an adaptable workflow to evaluate PKI probes in various cell types and experimental systems to support the discovery of new medicines.

The TFAP approach may also have broad ramifications for basic research. Finding specific PKI consensus signatures for kinases with diverse biological functions suggests the existence of such characteristic signatures for other kinases, including poorly characterized ‘dark kinases.’ Comparing their PKI consensus signatures to ‘illuminated’ kinome may help to elucidate the roles and functional relationship of dark kinases.

As the enabling technology, we used our multiplexed reporter system, the FACTORIAL. We showed previously that its homogeneous design renders it virtually impervious to multiple experimental variables, resulting in exceptional reproducibility and the signal/noise ratio(*14*). These factors permitted the accurate detection of comparatively weak TF responses to PKIs. Furthermore, we should underline that our approach entails the correlation analysis of TF constellations (i.e., TFAP signatures) rather than comparing individual TF responses. That was another essential factor contributing to reliable PKI evaluation.

It is important to notice that the TFAP approach does not compete with but complements existing target-based approaches. For example, one limitation of our approach stems from the point that inhibitors of different kinase family members have indistinguishable signatures. The reason for that is unclear. One possibility is that PKI consensus signatures reflect cells’ response to inhibited kinase families, regardless of specific mechanisms. That aligns with our previous observations that different perturbagens of the same biological system produced identical signatures, regardless of where and how they interfere(*12*). In support of that, CDK inhibitors exhibited the consensus signature that matched the consensus signature of DNA damage response, which is consistent with the key role of this kinase family in DNA damage response(*25*). Another possibility is that a limited number of TF endpoints restricts FACTORIAL’s resolution. If so, the resolution can be improved by expanding the FACTORIAL with additional TF reporters. Nonetheless, the exact PKI targets within the family can be pinpointed using a complementary target-based technique, such as cell-permeable energy transfer probes(*47*).

In summary, the TFAP approach illuminates PKI biology invisible to target-based approaches, thus expanding the toolbox for kinase research and PKI drug development.

## Materials and Methods

### Cells and reagents

The FACTORIAL assay was conducted in the HG19 subclone (Attagene) of the human hepatocyte HepG2 cell line [American Type Culture Collection (ATCC) #HB-8065]. This subclone was selected based on elevated xenobiotic metabolizing activity(*20*). Cells were maintained in a Dulbecco’s modified Eagle’s medium (DMEM) growth medium supplemented with 1% charcoal-stripped fetal bovine serum and antibiotics.

Kinase inhibitors were purchased from Tocris (www.tocris.com), Selleck Chemicals (www.selleckchem.com), and Cayman Chemical (www.caymanchem.com). To avoid the possible artifacts and ensure the reproducibility, we evaluated different batches of kinase inhibitors from multiple vendors. Inhibitors were dissolved in dimethyl sulfoxide (DMSO), keeping DMSO content in cell growth medium below 0.2%.

### Cell viability

Cell viability was evaluated by the XTT [2,3-bis-(2-methoxy-4-nitro-5-sulfophenyl)-2H-tetrazolium-5-carboxanilide] assay (ATCC) in HepG2/HG19 cells. As a baseline, we used cells treated with corresponding dilutions of the vehicle (DMSO).

### The mitochondria assay

We assessed PKIs’ effects on the mitochondria function using a galactose viability test in HepG2 cells(*48*). This assay is based on the observations that HepG2 cells can survive a complete mitochondrial inhibition utilizing glycolysis-generated ATP. Replacing media glucose with galactose inhibits ATP production by glycolysis, making these cells susceptible to mitochondrial toxicants(*48*).

Therefore, we compared HepG2 cells’ viability after incubation with tested compounds in glucose- or galactose-containing DMEM media. Specifically, cells were plated in 96-well plates at 10^4^ cells/well density. Twenty-four hours later, cells were rinsed twice with a glucose-free DMEM medium (Gibco, #11966-025) and incubated in a DMEM medium containing 10 mM glucose or galactose and 0.1% FBS. After a 24-h treatment with tested compounds, cell viability was assessed by XTT assay.

### The FACTORIAL® assay

The FACTORIAL assay in HepG2/HG19 cells was conducted as described(*12, 14*). The mix of 47 RTU plasmids was transiently cotransfected in a suspension of assay cells using TransIT-LT1 reagent (Mirus). The transfected cells were plated into 12-well plates, each well providing an independent FACTORIAL assay. Twenty-four hours later, cells were rinsed and incubated for another 24 hours with the evaluated compounds. Total RNA was isolated, and the RTU activity profiles were assessed by consecutive steps of RT-PCR amplification, Hpa I digest, and capillary electrophoresis, as described(*12, 14*) (Fig. S1).

### TFAP signatures

TF activity fold-changes were calculated by dividing the RTU activity values in compound-treated cells by those in vehicle-treated cells. The TFAP signatures are radial graphs with 47 axes showing the FACTORIAL RTUs’ fold changes on a logarithmic scale. The value of 1.0 indicates unchanged activity of corresponding TF.

The pair-wise similarity of TFAP signatures was assessed as the Pearson correlation coefficient ***r***. The signatures with the similarity r>0.70 were considered identical. In addition, we calculated the Euclidean distance ***d*** from the null signature to discriminate the experimental noise. The PKI signatures with ***d***<0.10 were considered the null signature. In those cases when PKI treatment caused cell death, RNA degradation resulted in ‘empty’ signatures.

### The PKI consensus signatures

To obtain the PKI consensus signature for a given kinase family, we clustered PKIs’ TFAP signatures at multiple concentrations using a recurrent agglomerative hierarchical clustering algorithm(*12*). The cluster with most PKI signatures was regarded the main cluster, and the center cluster signature was presumed the PKI consensus signature. The similarity of individual PKI signatures and the consensus signature exceeded the identity threshold (*r*>0.70).

## Acknowledgements

We thank D. Drewry, L.Graves, W. Zuercher, R. Medzhitov, and A. Gudkov for critique and helpful suggestions. Funding: This work was supported by NIH grants R44GM125469 and R44GM136083.

## Author contributions

A.M. and S.S.M. designed the research. A.M., L.M., E.M., K.L.G., and B.L. performed the research. S.M. Jr. developed algorithms for data analysis. A.M. and S.S.M. analyzed the data. S.S.M. wrote the manuscript with input from A.M.

## Competing interests

A.M., L.M., E.M., and S.S.M. have competing financial interests as Attagene employees and shareholders. A.M. and S.S.M. are inventors on a U.S. patent related to this work (no. 7,700,284, issued on 20 April 2010). The authors declare no other competing interests.

## Data and materials availability

All data needed to evaluate the papers’ conclusions are present in the paper and/or the Supplementary Materials. Additional data related to this paper may be requested from A.M. or S.S.M.

## Supplementary Materials

**Fig. S1.**
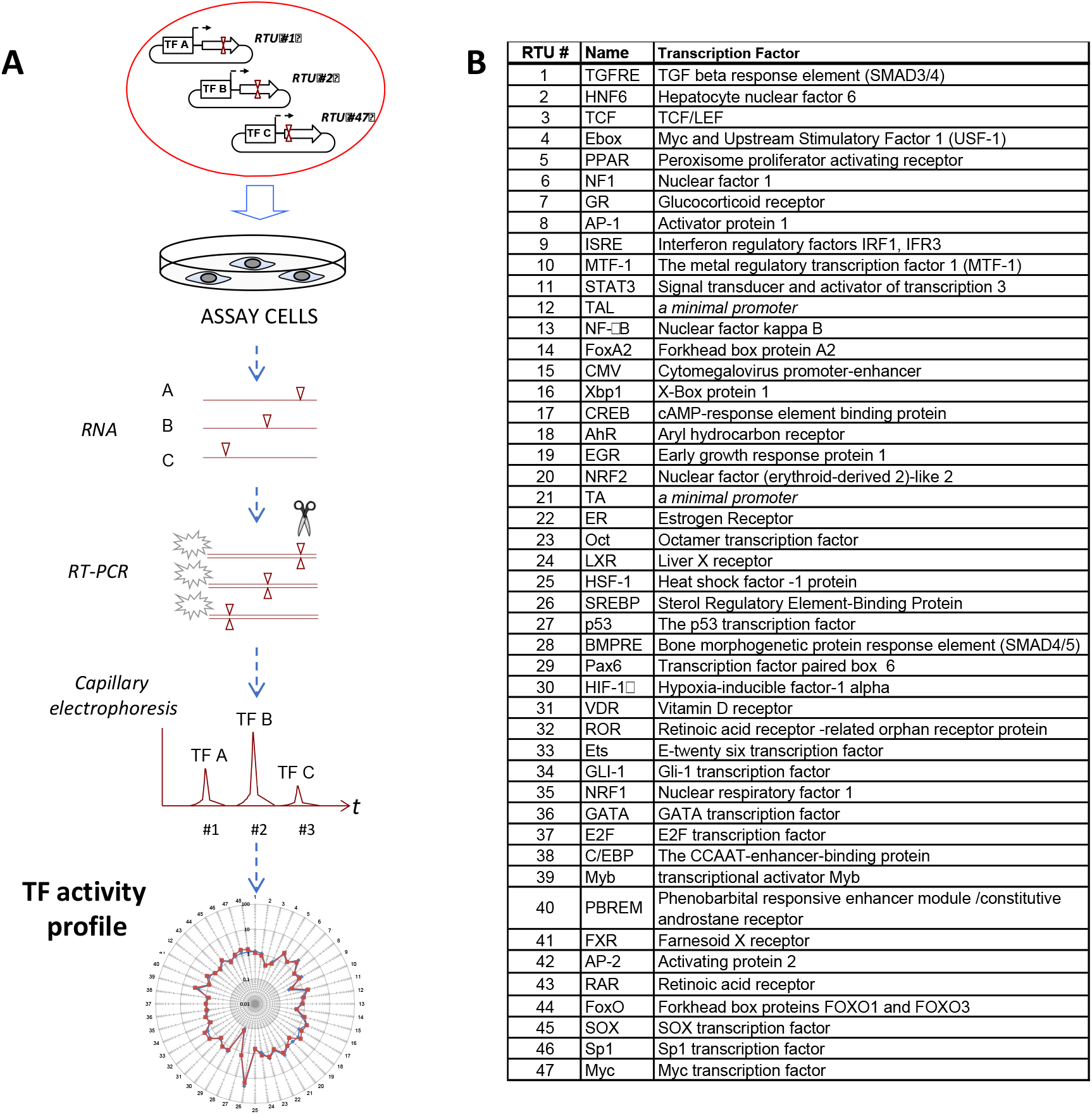
The workflow and endpoints of the FACTORIAL assay.

**Fig. S2.**
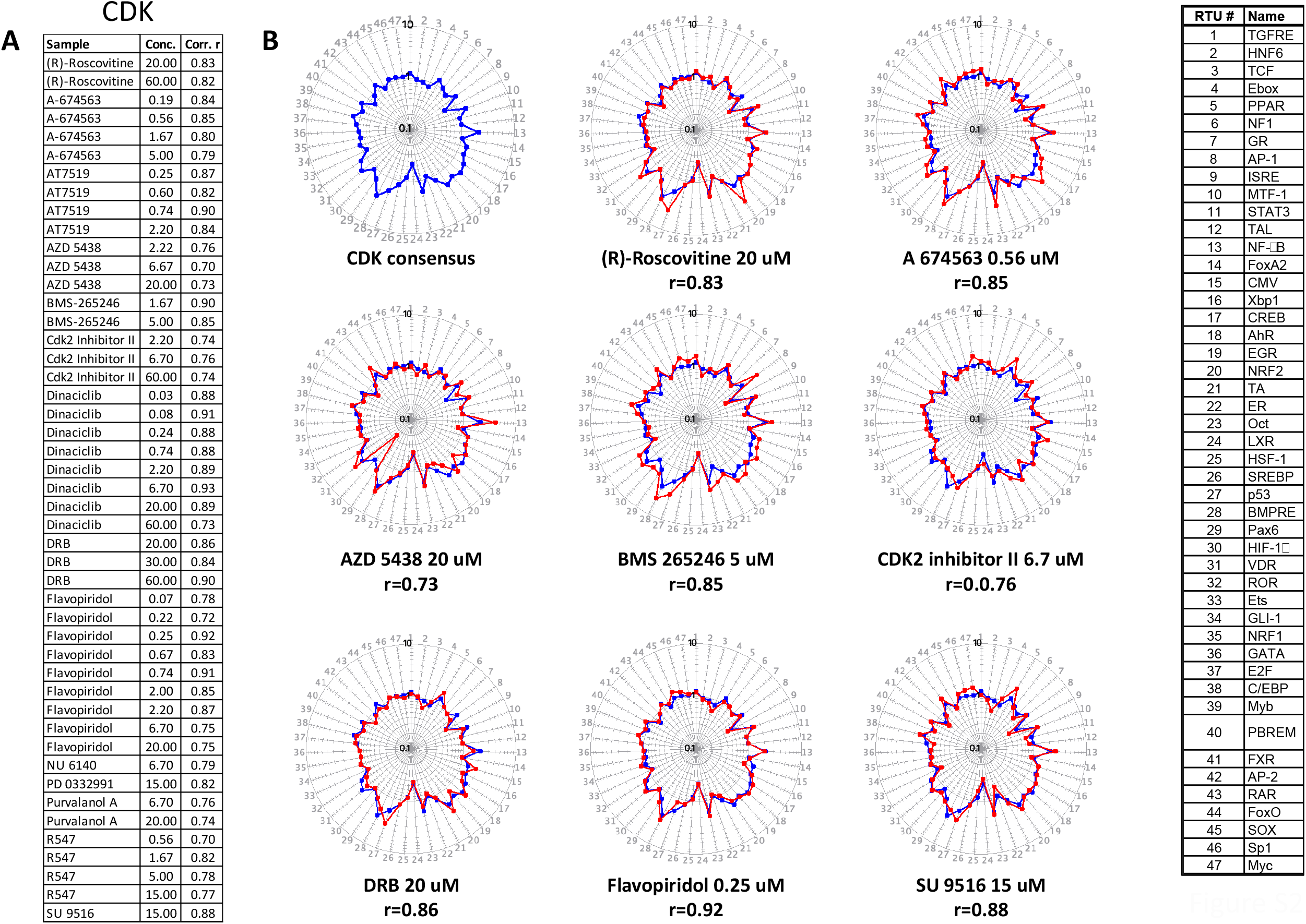
The TFAP signatures of CDK inhibitors.

**Fig. S3.**
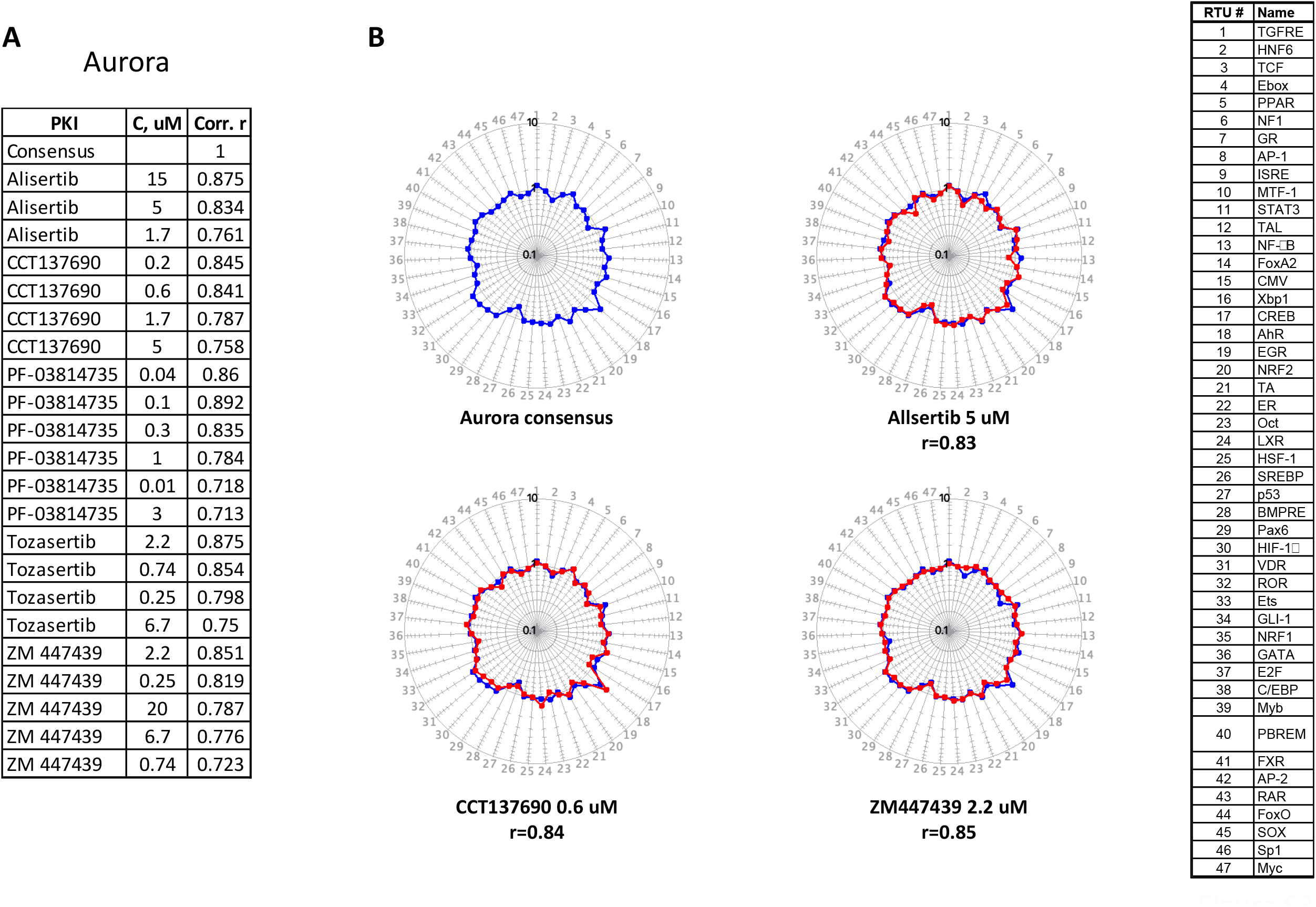
The TFAP signatures of CDK inhibitors.

**Fig. S4.**
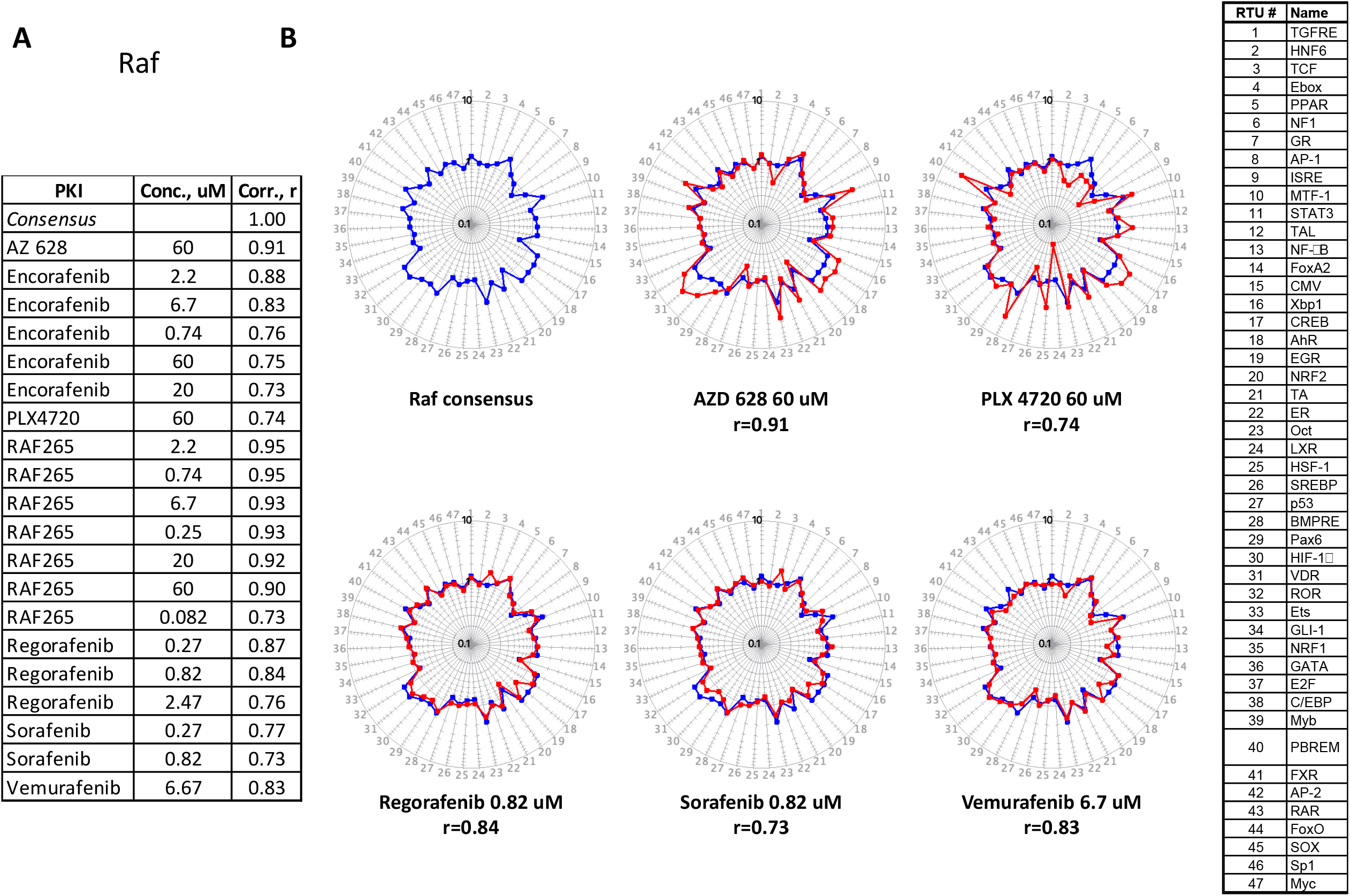
The TFAP signatures of RAF inhibitors.

**Fig. S5.**
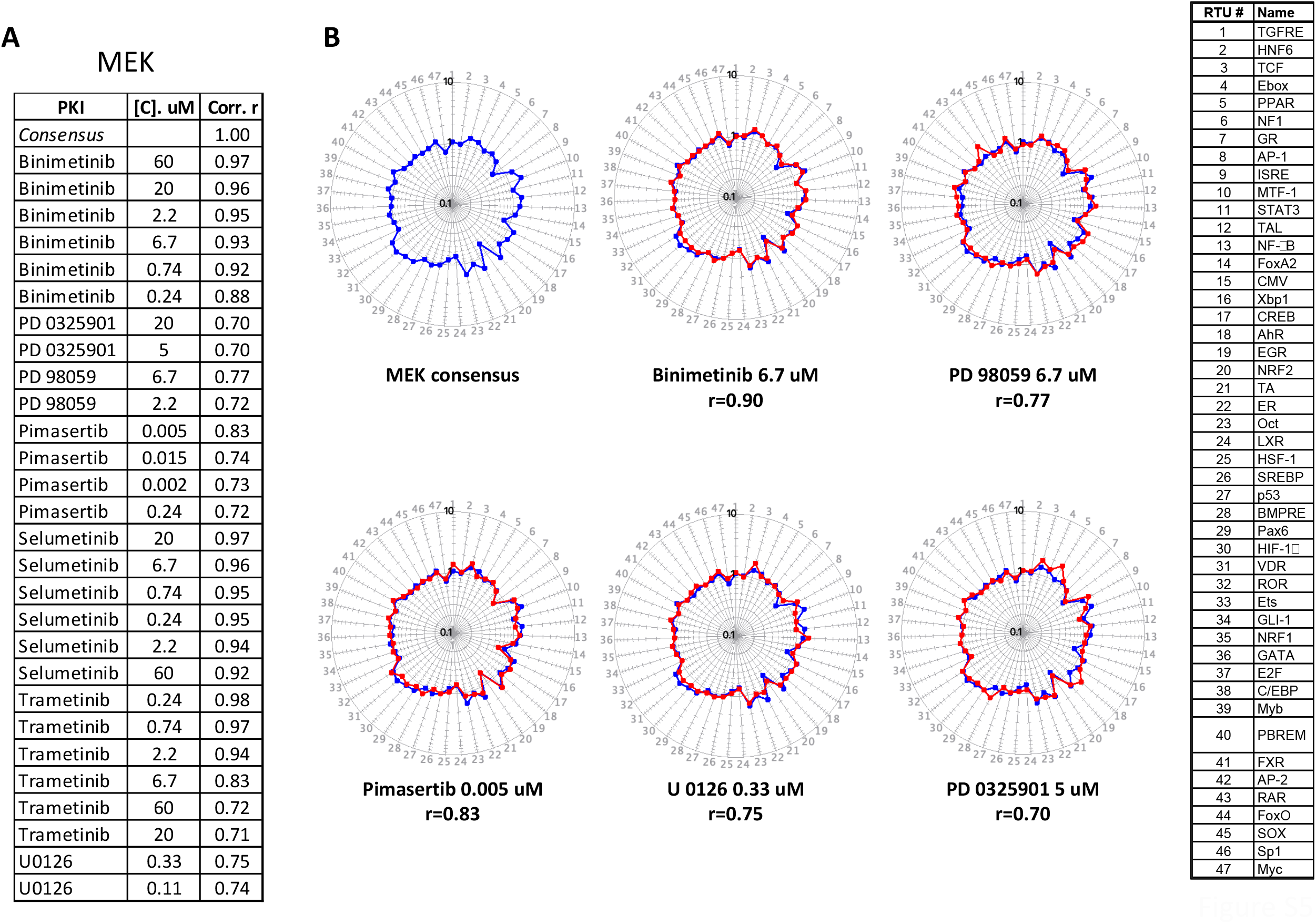
The TFAP signatures of MEK inhibitors.

**Fig. S6.**
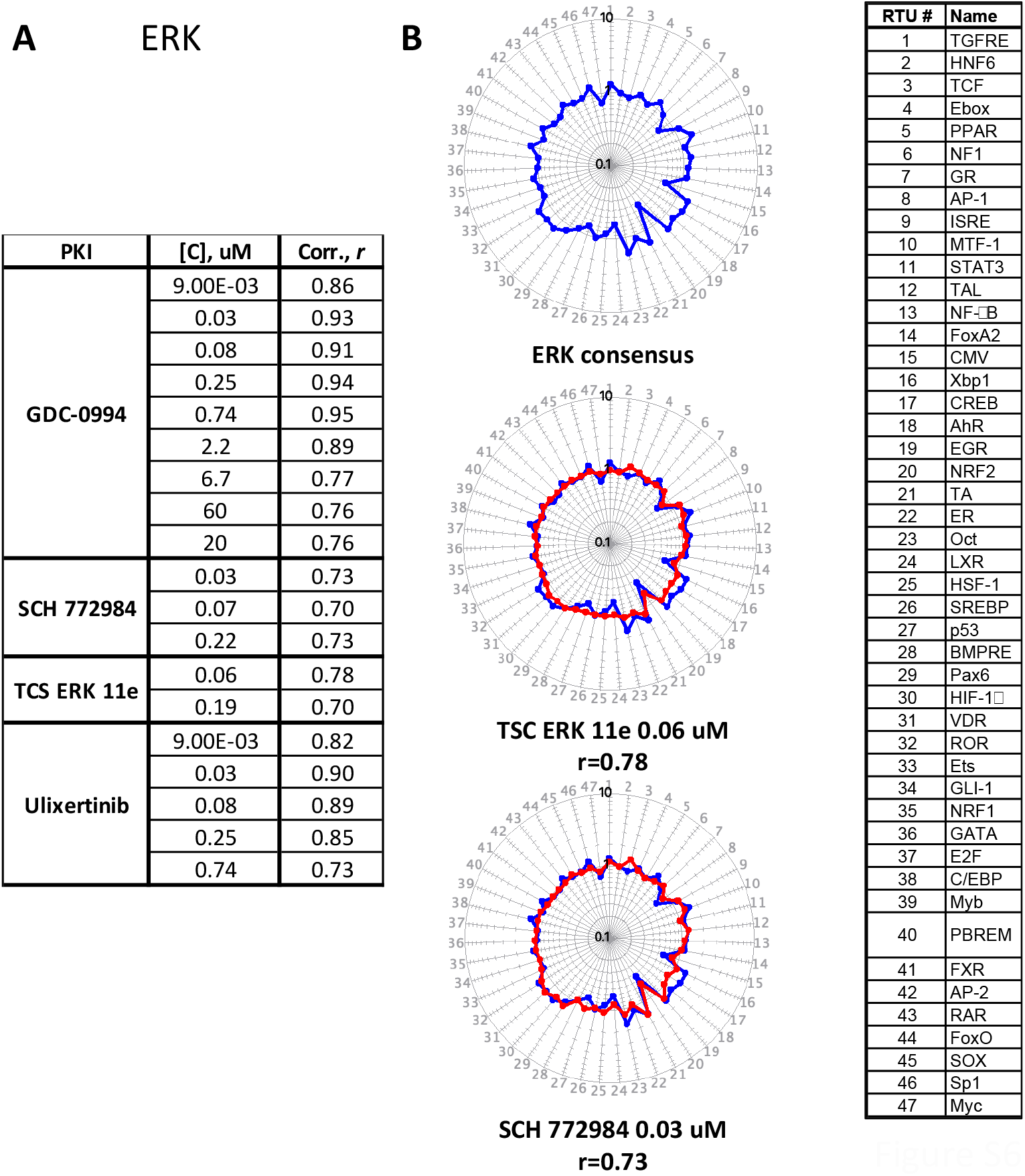
The TFAP signatures of ERK inhibitors.

**Fig. S7.**
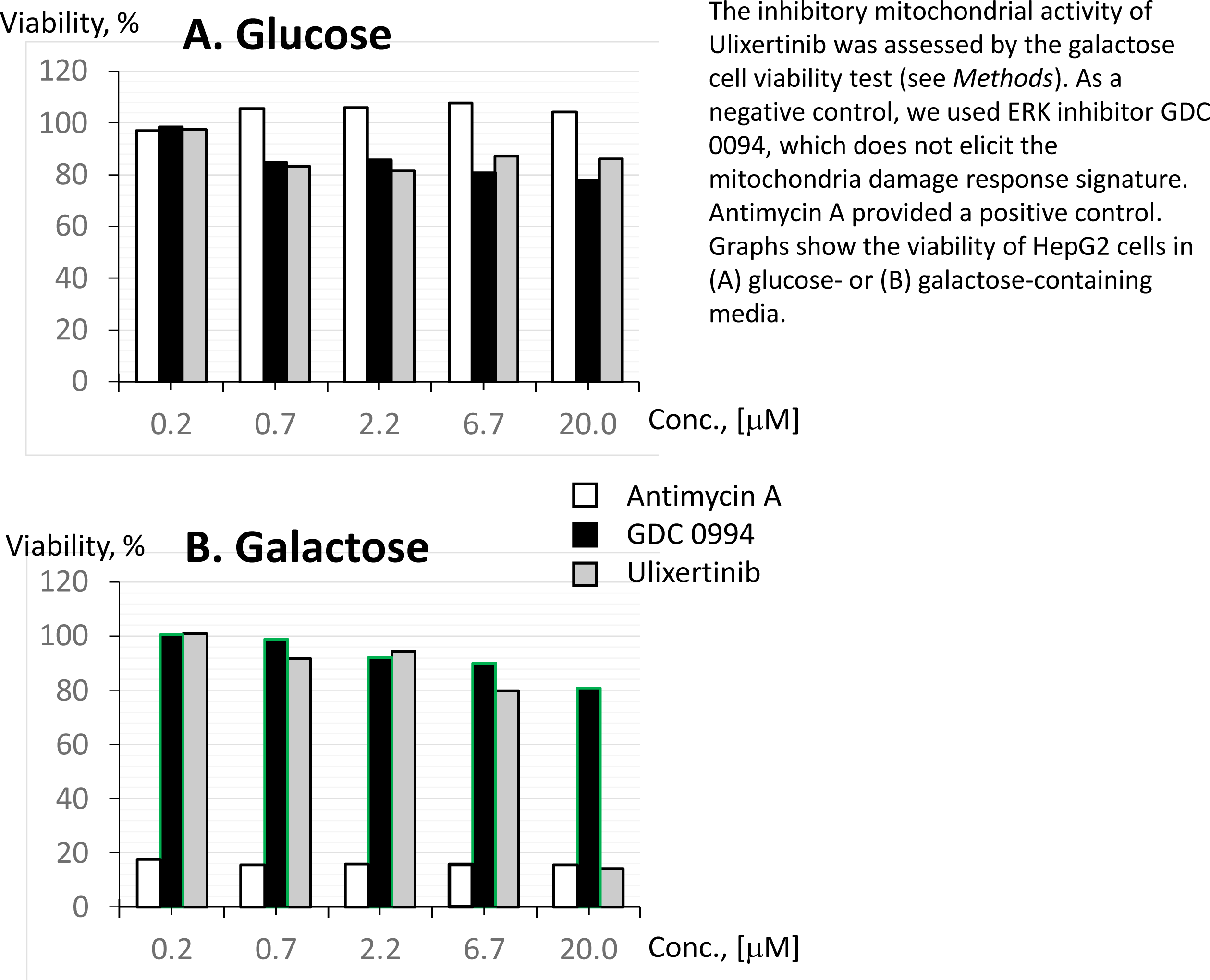
Mitochondria inhibition by kinase inhibitors.

## Notes

### Summary of Updates

General revision for the text clarity.

